# Fabry Cardiomyopathy: Myocardial Fibrosis, Inflammation and Down-Regulation of Mannose6Phosphate Receptors cause Low accessibility to Enzyme Replacement Therapy

**DOI:** 10.1101/2023.12.20.572710

**Authors:** Andrea Frustaci, Romina Verardo, Michele Magnocavallo, Emanuela Frustaci, Matteo Antonio Russo, Cristina Chimenti

## Abstract

**Background:** Clinical impact of enzyme replacement therapy (ERT) on advanced Fabry disease cardiomyopathy (FDCM) appears limited. The pathologic mechanisms involved are still unclear.

**Methods:** Ten male patients with advanced FDCM (echocardiographic maximal wall thickness 19.3 ± 2.1 mm) underwent left ventricular endomyocardial biopsy before and 4 hours after beta-agalsidase infusion (1 mg/Kg). Comparative studies between pre and post infusion samples included: histology and electron microscopy; assessment of myocardial alpha-galactosidase A activity; immunohistochemistry for alpha-galactosidase A and semiquantitative evaluation (from 0 to 3) of its cardiomyocyte content; Ultrastructural immunogold analysis with anti-alpha-galactosidase A ab. Western Blot (WB) quantification of mannose-6-phosphate receptors (M6Pr). Controls were surgical left ventricular biopsies from patients with mitral stenosis.

**Results:** Histologic and Ultrastructural evaluation showed no removal of storage material while myocardial fibrosis was 9.8 ± 6.8 vs 3.8 ± 2.0 of controls and virus-negative lymphocytic inflammation was observed in 7 out of 10 patients. At Ultrastructural immunogold analysis, Myocardial alpha-galactosidase A activity increased in post infusion samples by overall 1.89-fold. Alpha-galactosidase A immunostaining in cardiomyocytes was absent at baseline in all patients and did not significantly improve in post-infusion samples. Immunogold particles increased by 1.33**-**fold (17.6 ± 3.6 pre infusion vs 21.5 ± 5.9 post) remaining far from normal controls (86.9 ± 6.6). Protein analysis showed M6Pr in advanced FDCM to be 81% lower than in normal heart.

**Conclusions:** Our study shows a low accessibility to ERT of cardiomyocytes affected by advanced FDCM. It is sustained by myocardial fibrosis, inflammation and severe down-regulation of M6Pr.

## Introduction

Fabry disease (FD) is an X-linked disorder caused by deficiency of lysosomal enzyme alpha galactosidase A, resulting in progressive intracellular accumulation of glycosphingolipids, essentially globotriacylceramide (GB3) in different tissues, including skin, kidneys, vascular endothelium, ganglion cells of peripheral nervous system and heart ^1^. Cardiac involvement is characterized by progressive left ventricular hypertrophy that mimic the morphological and clinical features of hypertrophic cardiomyopathy ^2–4^ and is the most important cause of death in both hemizygote males and heterozygote females ^1^.

After more than 20 years from instauration of enzyme replacement therapy (ERT) for FD, cardiac clinical results are still controversial as a reduction of left ventricular hypertrophy has been recognized in some studies ^5,6^, confirmed only in those patients with less advanced disease in other reports ^7^ and denied in others ^8,9^. On the other hand, a minimal reduction of myocardial mass doesn’t necessarily correspond to improvement of cardiac infiltration but, particularly for the most advanced phases, can be the result of undesirable events as cardiac cell death and fibrosis ^10^. In addition, post-mortem studies reported persistence of myocardial infiltration after 2.5 years of ERT ^11^, while recent experimental studies raise doubts on the ability of the infused enzyme to enter affected cardiomyocytes and remove glycosphingolipids accumulation from lysosomes ^12^. Other studies showed that, even if the enzyme enter the cardiomyocytes, it is unable to remove the long-lasting stored material ^13^. Pathologic as well as molecular mechanisms that limit ERT impact particularly in the advanced phase of FD cardiomyopathy (FDCM) are still unclear.

In the present study we evaluated the ability of ERT to increase enzyme activity in the myocardium, to enter the cardiomyocytes and to remove the storage material, performing left ventricular endomyocardial biopsy (EMB) before and 4 hours after ERT infusion in FD patients with severe left ventricular hypertrophy. Cardiomyocyte accessibility to ERT was compared with histologic and ultrastructural findings and with myocardial expression of mannose-6-phosphate receptors (M6Pr) which represent the main molecular pathway for ERT uptake and internalization into cardiac cells.

## Methods

### Patient population and clinical studies

Ten male patients with advanced FDCM constituted our population. For all patients, reduced peripheral blood alpha-galactosidase A activity was detected as previously described ^14^ and causal mutations were identified by direct sequencing of alpha-galactosidase A gene in all families. Extensive clinical examination, including the assessment of FD systemic manifestations, ECG, 2D echocardiography with Doppler analysis were performed in all patients ^15^. Maximal wall thickness was defined as the greatest thickness measured at any segment of left ventricular wall.

### Cardiac catheterization and endomyocardial biopsy

Invasive studies were performed after patient written informed consent and approval by the Ethical Committees of our Institution and included cardiac catheterization, coronary angiography, left ventricular angiography and EMB. Four to six biopsy samples were drawn before and 4 hours after beta-agalsidase infusion at standard dosage (1 mg/kg). Comparative studies between pre and post infusion samples were performed and included: a) histology, b) transmission electron microscopy (TEM), c) immunohistochemistry for alpha-galactosidase A, d) Ultrastructural immune-gold analysis with anti-alpha galactosidase A Ab. e) In pre-infusion frozen samples it was obtained a Western blot quantification of Mannose-6-Phosphate receptors (M6Pr). Controls were 10 surgical left ventricular biopsies from male patients with mitral stenosis and normal left ventricular function, undergoing surgical valve replacement.

### Histological, morphometric and molecular studies

Myocardial samples were processed for histological and histochemical analyses, and for TEM as previously described ^15,16^.

Histologic sections stained with Masson’s trichrome were examined at 400x magnification with a reticule containing 42 sampling points (105844, Wild Heerbrugg Instruments, Gals, Switzerland) to determine the percent area occupied by cardiomyocytes, by interstitial and replacement fibrosis and by other components (vascular spaces, interstitial cells) ^15^.

Morphometric analysis of percent of vacuolar area occupied by myelin figures was performed on 10 photographic negatives of TEM sections/each patient corresponding to 30±7 (range 20-40) transversally cut cardiomyocytes.

Histologic diagnosis of myocarditis included evidence of leukocyte infiltrates in association with damage of the adjacent myocytes, according to the Dallas criteria ^17^ confirmed by immunohistochemistry ^18^. In particular for the phenotypic characterization of the inflammatory infiltrates immunohistochemistry for CD3, CD20, CD43, CD45RO and CD68 was performed (Dako, Carpinteria, California, USA). The presence of an inflammatory infiltrate ≥14 leucocytes/mm2 including up to 4 monocytes/mm2, with the presence of CD3-positive T-lymphocytes ≥7 cells/mm2 associated with evidence of degeneration and/or necrosis of the adjacent cardiomyocytes, was considered diagnostic for myocarditis ^18^.

In patients with histologic evidence of overlapping myocardial inflammation, Real Time Pcr for the most common cardiotropic viruses (Adenovirus, Epstein Barr Virus, Herpes virus, Parvovirus B19, Cytomegalovirus, Enterovirus, Influenza A/B) was obtained in 2 frozen endomyocardial samples.

### Alpha-galactosidase A activity and Immunohistochemistry

Myocardial alpha-galactosidase A activity was assessed on preinfusion and postinfusion frozen samples as previously described ^14^. Briefly, the EMB specimens were homogenized in 1 mM citrate–phosphate buffer (pH 4.5) and then centrifuged. The resulting endomyocardial supernatants were assayed with 4-methylumbelliferyl-alpha*-*D-galactoside for alpha galactosidase A activity (measured as nanomoles per hour per milligram of protein).

Immunohistochemistry for alpha-galactosidase A was performed on 5μm cut frozen sections using a rabbit polyclonal antibody (courtesy from Genzyme Corp., Cambridge, Massachusetts, USA) Tissue sections were fixed for 5 minutes with 10% neutral buffered formalin, exposed to standard blockade of endogenous peroxidase (Dako Denmark A/S) for 15 minutes and pretreated by microwaving in 10mM citrate buffer, pH 6.0 for 10 minutes. The specimens were then incubated with PBS containing 10% of normal goat serum for 30 minutes at RT to block non-specific binding and subsequently rabbit anti-human alpha-galactosidase A antibody at a dilution of 1:100 was applied to tissue sections. Sections were incubated overnight at 4°C, then washed in PBS and exposed to the detection kit (EnVision + System-HRP Anti-Rabbit, Dako Denmark A/S) which included 3,3′-diaminobenzidine tetrahydrochloride as chromogen and counterstained with Harris hematoxylin. Negative controls were sections of FD patients stained without using the primary antibody and slides from normal controls. In each biopsy, the intensity of the immunostaining for alpha-galactosidase A in cardiomyocytes was graded semiquantitatively as follows: score 0=absent; score 1=mild; score 2=moderate; score 3=marked.

### Electron microscopy

For immunogold analysis small fragments of cardiac biopsies were fixed in 2% freshly depolymerised paraformaldehyde and 0.2% glutaraldehyde in 0.1M cacodilate buffer pH 7.4 for 1 hour at 4°C. Samples were rinsed in the same buffer, partially dehydrated and embedded in London Resin White (LR White, Agar Scientific Ltd). Ultrathin sections were processed for immunogold technique. Grids were pre-incubated with 10% normal goat serum in 10 mM PBS containing 1% bovine serum albumine and 0.13% NaN3 (medium A), for 15 minutes at RT; sections were then incubated with primary antibody, rabbit polyclonal anti-alpha-galactosidase (sc-25823 Santa Cruz Biotechnology, INC) diluted 1:100 in medium A for 1 hour at RT. After rinsing in medium A containing 0.01% Tween 20 (Merck), sections were incubated in goat anti-rabbit IgG conjugated to 15 nm colloidal gold (British BioCell Int.), diluted 1:30 in medium A, containing fish gelatine, for 1 hour at RT. Grids were thoroughly rinsed in distilled water, contrasted with aqueous 2% uranyl acetate for 20 min. and photographed in a Zeiss EM 900 electron microscope.

### Assessment of myocardial Mannose-6-phosphate receptors

We determined the expression of M6Pr in frozen myocardial tissue. Results were compared with values from surgical control unloaded myocardium (papillary muscle of patients with mitral stenosis undergoing valve replacement). The expression of M6Pr, molecular weight 275 kDa, was visualized by using IGF-II Receptor/CI-M6Pr, (D8Z3J) polyclonal antibody (1:1000, Cell Signaling Technolo-gy). Anti-α-sarcomeric actin antibody (1:500, Sigma-Aldrich), molecular weight 43 kDa, was used for normalization. Signal was visualized using a secondary horseradish-peroxidase-labeled goat anti-mouse antibody (goat anti-mouse IgG-HRP 1:5000, SantaCruzBiotechnology) and enhanced chemiluminescence (ECL Clarity Bio-Rad). The purity as well as equal loading (40γ) of the protein was determined by Nanodrop One (Thermofisher). To normalize target protein expression, the band intensity of each sample is determined by densitometry with Image J software. Next, the intensity of the target pro-tein is divided by the intensity of the loading control protein.

This calculation adjusts the expression of the protein of interest to a common scale and reduces the impact of sample-to-sample variation. Relative target protein expression can then be compared across all lanes to assess changes in target protein expression across samples. Digital images of the resulting bands were quantified by the Image Lab software package (Bio-Rad Labora-tories, Munchen, Germany) and expressed as arbitrary densitometric units.

### Statistical analysis

Descriptive statistics were used to summarize the data; results are reported as medians and interquartile ranges or means and standard deviations, as appropriate. Categorical variables were summarized as counts and percentages. Baseline demographic and clinical characteristics will be presented in table format. Changes of variables before and after 4 hours of ERT infusion were analyzed with paired *t* test or Signed Rank test in case of non normality. A two tailed p value ≤ 0.05 was considered statistically significant. All statistical analyses were performed using R statistical analysis software.

## Results

### Patient population

We enrolled ten male patients (mean age: 45.0±7.3 years) with advanced FDCM (maximal wall thickness 19.3±2.1 mm); all of them were on treatment with beta-agalsidase (1mg/Kg/every other week). All clinical, echocardiographic and genetic characteristics of the population are shown in *Table 1* and *Table 2*. Patient 4 and 8 were affected by the cardiac variant of FD, carrying the same gene mutation but pertaining to different families.

**Table 1.** Characteristics and response to enzyme replacement therapy of 10 male patients with advanced Fabry Disease Cardiomyopathy. * S= skin, CNS=central nervous system; E= eyes, EA=ears, H=hypoacusia, AP=acroparesthesias; K= kidney. † MWT= Maximal wall thickness; EF=Ejection fraction. § nmol/h/mg of Protein, values are the mean (± SD). Lymphocytic values are the results of three independent determinations on peripheral blood lymphocytes, myocardial values were determined on two samples before and two samples after infusion; ‡ Semiquantitative evaluation of immunostaining graded as follows: score 0=absent; score 1=mild; score 2=moderate; score 3=marked; ||Calculated as percent of vacuolar area occupied by myelin figures on 10 transmission electron microscopy photographic negatives/each patient.

| Patient Population |  |  |  |  |  |  |  |  |  |  |
| --- | --- | --- | --- | --- | --- | --- | --- | --- | --- | --- |
| Characteristics | 1 | 2 | 3 | 4 | 5 | 6 | 7 | 8 | 9 | 10 |
| Age, ys | 40 | 42 | 43 | 53 | 31 | 44 | 53 | 52 | 52 | 40 |
| Genotype | Q279K | G328R | c946del | N215S | C378Y | R112C | Y216C | N215S | c427del | R220X |
|  |  |  | G |  |  |  |  |  | ACTT |  |
| Extracardiac | S, CNS, | E, K | S, CNS, | None | S, E, AP | S, CNS, | S, E, H, | None | AP, K, | S, E, EA, |
| symptoms* | E, EA, K |  | E EA, H, |  |  | E, EA, | K |  | H,S | K |
|  |  |  | AP, K |  |  | H, AP, K |  |  |  |  |
| Ecocardiography† |  |  |  |  |  |  |  |  |  |  |
| MWT, mm | 18.5 | 20.0 | 21.0 | 16.5 | 20.0 | 17.0 | 23.0 | 19.0 | 21.0 | 17.0 |
| EF, % | 65 | 58 | 57 | 60 | 50 | 56 | 67 | 65 | 55 | 59 |
| *Enzymatic activity§* |  |  |  |  |  |  |  |  |  |  |
| Lymphocytic | 15.2 | 141.0 | 20.9 | 150.7 | 103.1 | 30.6 | 79.2 | 50.2 | 19.7 | 23.1 |
| at diagnosis | ±0.8 | ±5.6 | ±1.5 | ±7.2 | ±3.5 | ±6.3 | ±84.5 | ±5.6 | ±1.3 | ±2.6 |
| Myocardial | 83.3 | 52.6 | 85.4 | 98.2 | 105.1 | 59.3 | 148.6 | 81.2 | 84.1 | 118.0 |
| before infusion | ±4.3 | ±3.1 | ±3.0 | ±2.6 | ±6.9 | ±1.1 | ±7.6 | ±1.6 | ±1.3 | ±4.0 |
| Myocardial | 119.2 | 123.5 | 172.9 | 200.8 | 177.6 | 121.2 | 222.9 | 182.0 | 151.6 | 264.9 |
| after infusion | ±9.4 | ±4.9 | ±10.0 | ±3.2 | ±14.1 | ±14.7 | ±16.8 | ±7.8 | ±11.9 | ±14.3 |
| *IHC for alphaGal‡* |  |  |  |  |  |  |  |  |  |  |
| Before infusion | 0 | 0 | 0 | 0 | 0 | 0 | 0 | 0 | 0 | 0 |
| After infusion | 1 | 0 | 0 | 1 | 0 | 1 | 0 | 1 | 0 | 1 |
| *Lysosomal GB3 (%)\|\|* |  |  |  |  |  |  |  |  |  |  |
| Before infusion | 50.0 | 49.3 | 53.3 | 48.5 | 44.8 | 49.9 | 45.2 | 51.7 | 55.9 | 43.3 |
| After infusion | 50.3 | 49.5 | 53.2 | 48.4 | 45.4 | 49.8 | 45.5 | 51.7 | 55.8 | 43.8 |

**Table 2.**
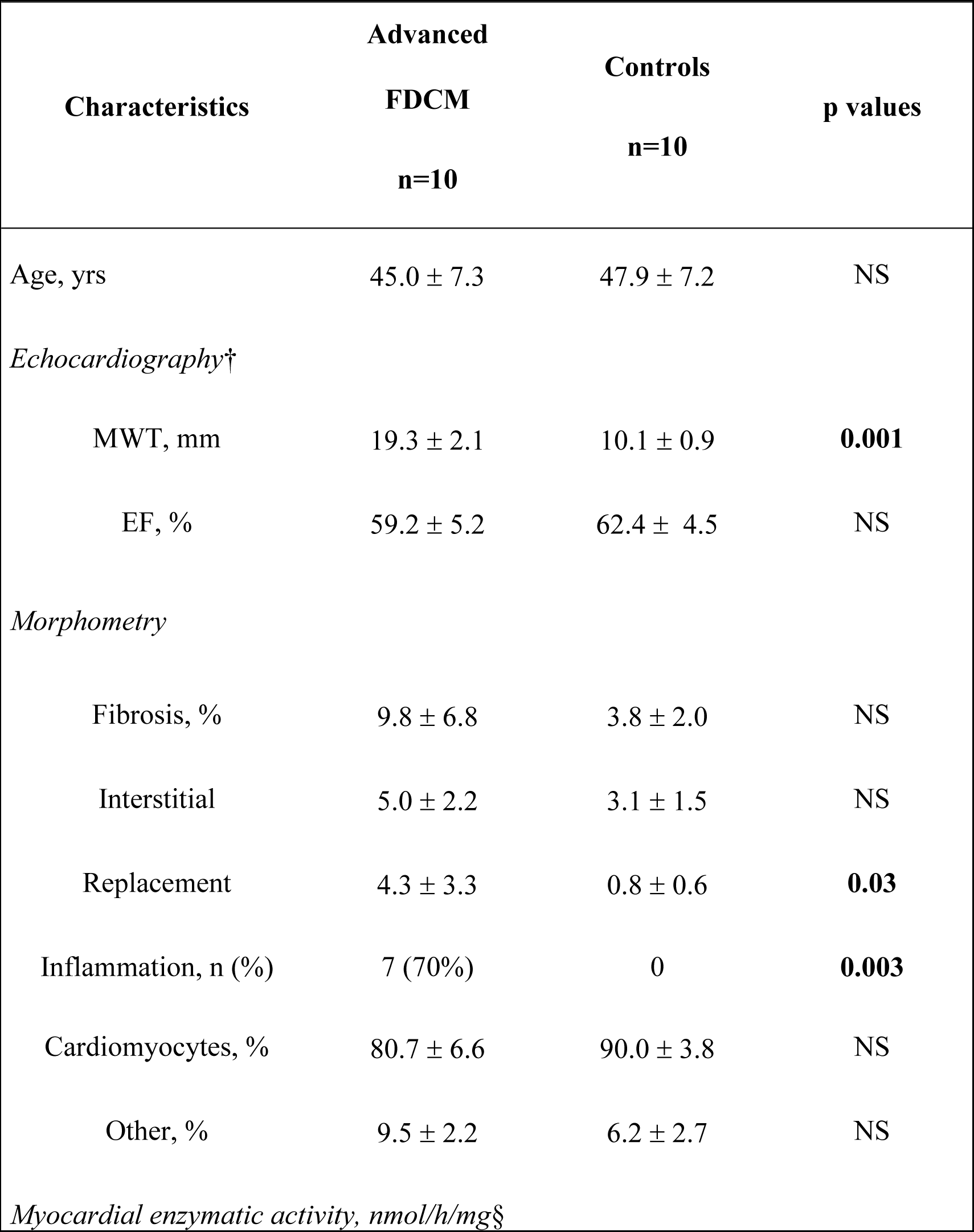
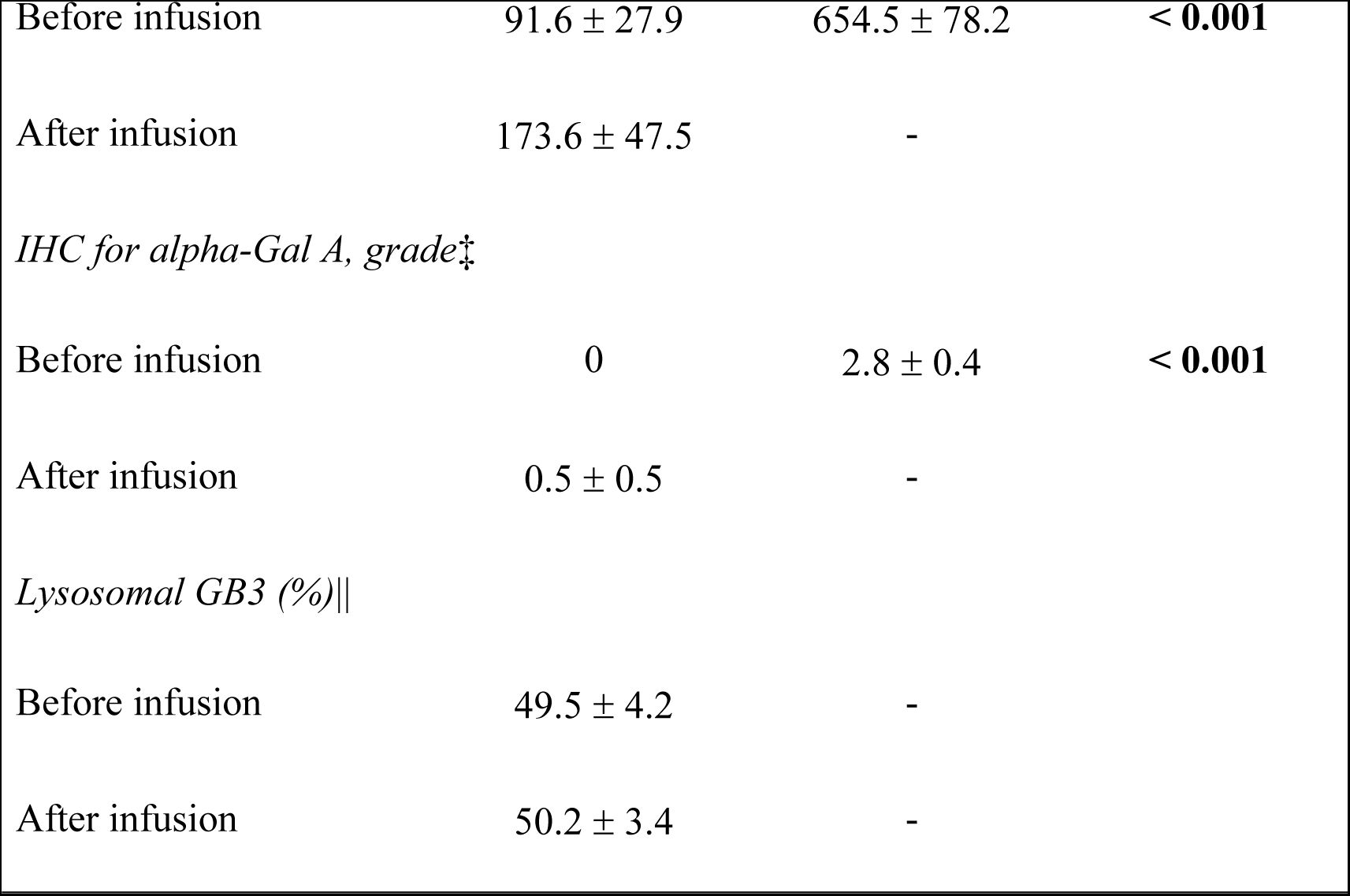
Comparison among echocardiographic, morphometric and enzymatic data of male patients with advanced Fabry Disease Cardiomyopathy and male controls. NS=Not statistically significant differences among groups. † MWT= Maximal wall thickness; EF=Ejection fraction. § Values are the results of enzymatic activity determination on two samples before and two samples after infusion. ‡ Semiquantitative evaluation of immunostaining graded as follows: score 0=absent; score 1=mild; score 2=moderate; score 3=marked. ||Calculated as percent of vacuolar area occupied by myelin figures on 10 transmission electron microscopy photographic negatives/each patient.

| Characteristics | Advanced<br>FDCM<br>n=10 | Controls<br>n=10 | p values |
| --- | --- | --- | --- |
| Age, yrs | 45.0 ± 7.3 | 47.9 ± 7.2 | NS |
| *Echocardiography*† |  |  |  |
| MWT, mm | 19.3 ± 2.1 | 10.1 ± 0.9 | **0.001** |
| EF, % | 59.2 ± 5.2 | 62.4 ± 4.5 | NS |
| *Morphometry* |  |  |  |
| Fibrosis, % | 9.8 ± 6.8 | 3.8 ± 2.0 | NS |
| Interstitial | 5.0 ± 2.2 | 3.1 ± 1.5 | NS |
| Replacement | 4.3 ± 3.3 | 0.8 ± 0.6 | **0.03** |
| Inflammation, n (%) | 7 (70%) | 0 | **0.003** |
| Cardiomyocytes, % | 80.7 ± 6.6 | 90.0 ± 3.8 | NS |
| Other, % | 9.5 ± 2.2 | 6.2 ± 2.7 | NS |
| *Myocardial enzymatic activity, nmol/h/mg*§ |  |  |  |
| Before infusion | 91.6 ± 27.9 | 654.5 ± 78.2 | < **0.001** |
| After infusion | 173.6 ± 47.5 | - |  |
| *IHC for alpha-Gal A, grade<sup>‡</sup>* |  |  |  |
| Before infusion | 0 | 2.8 ± 0.4 | < **0.001** |
| After infusion | 0.5 ± 0.5 | - |  |

*Lysosomal GB3 (%)||*
|  |  |  |
| --- | --- | --- |
| Before infusion | 49.5 ± 4.2 | - |
| After infusion | 50.2 ± 3.4 | - |

### Histological, morphometric and molecular studies

In all FD patients histology showed the presence of large perinuclear vacuoles in cardiomyocytes, containing material that on frozen section stained positively with PAS and Sudan-Black, suggesting the accumulation of glycosphingolipids. These vacuoles consisted at electron microscopy of concentric lamellar figures in single-membrane bound vesicles (myelin bodies), diagnostic for FD (*Figures 1, 2 and 3*). Morphometric analysis showed a 3 fold increase in interstitial and replacement fibrosis in FD patients with advanced disease compared with controls (*Figure 1 and Table 2*). Ultrastructural examination in the former group showed that widening of the interstitium was due not only to fibrosis but also to extracellular storage material deriving from cell secretion and death (*Figure 1*). Virus-negative lymphocytic (CD45Ro positive) myocarditis was observed in 7 out of 10 patients enrolled in the study (*Figure 1 - Panel B*). Morphometry on electron micrographs showed in patients with advanced FDCM no removal of storage material from lysosomes (*Figure 1 - Panel C-D*).

**Figure 1.**
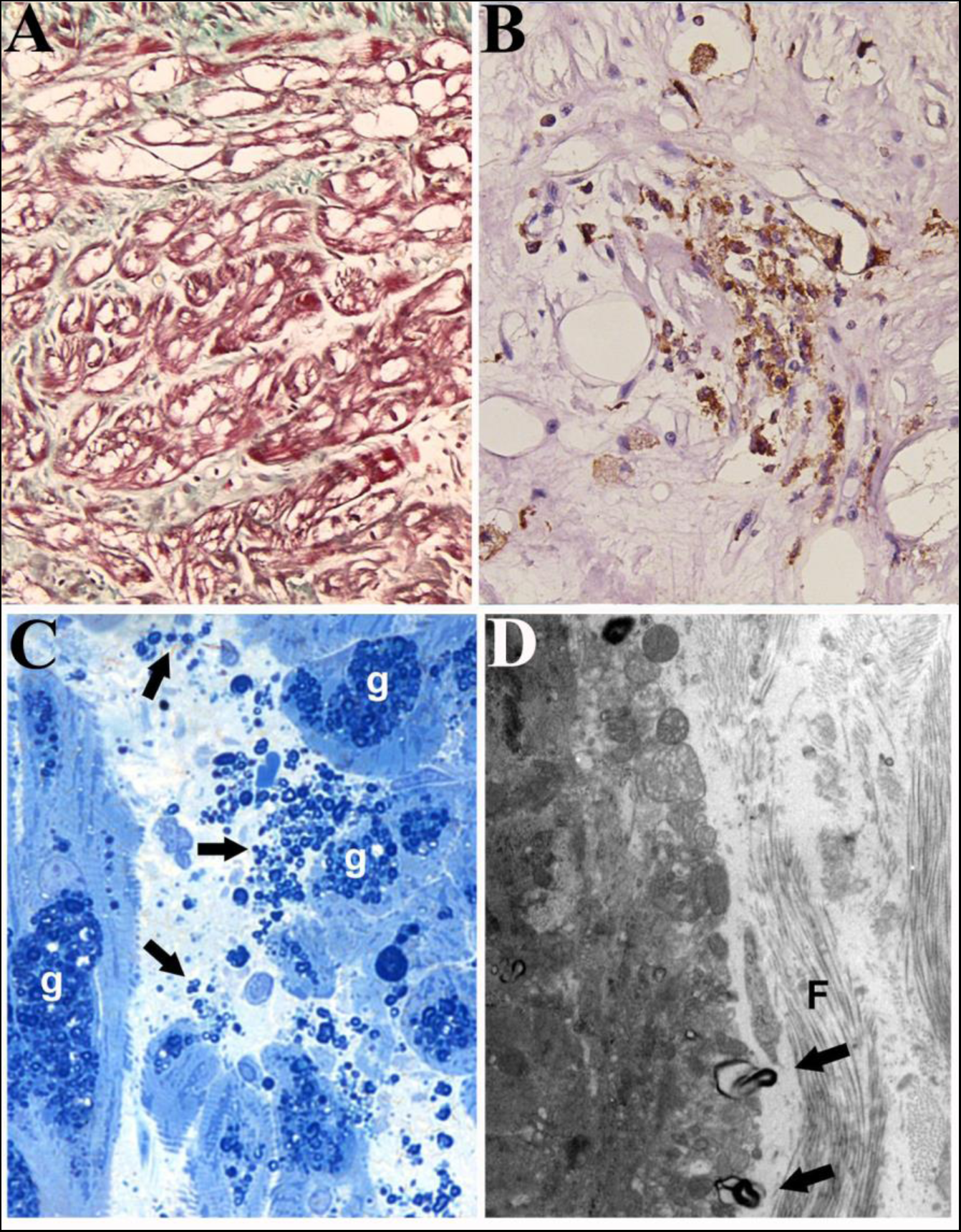
Histologic and ultrastructural characteristics of patients with severe Fabry Disease Cardiomyopathy. Panel A shows severe cellular glycolipid infiltration associated with diffuse interstitial fibrosis (Masson trichrome, 100x). Panel B reveals an overlapping lymphocytic CD45Ro+ myocardial inflammation with focal necrosis of adjacent cardiomyocytes. Panels C and D show Extracellular glycosphingolipids release (arrows, semithin sections, Azur II staining, 400x, G=glycosphingolipids) and secretion (arrows, transmission electron microscopy, uranyl acetate, 1600x).

**Figure 2.**
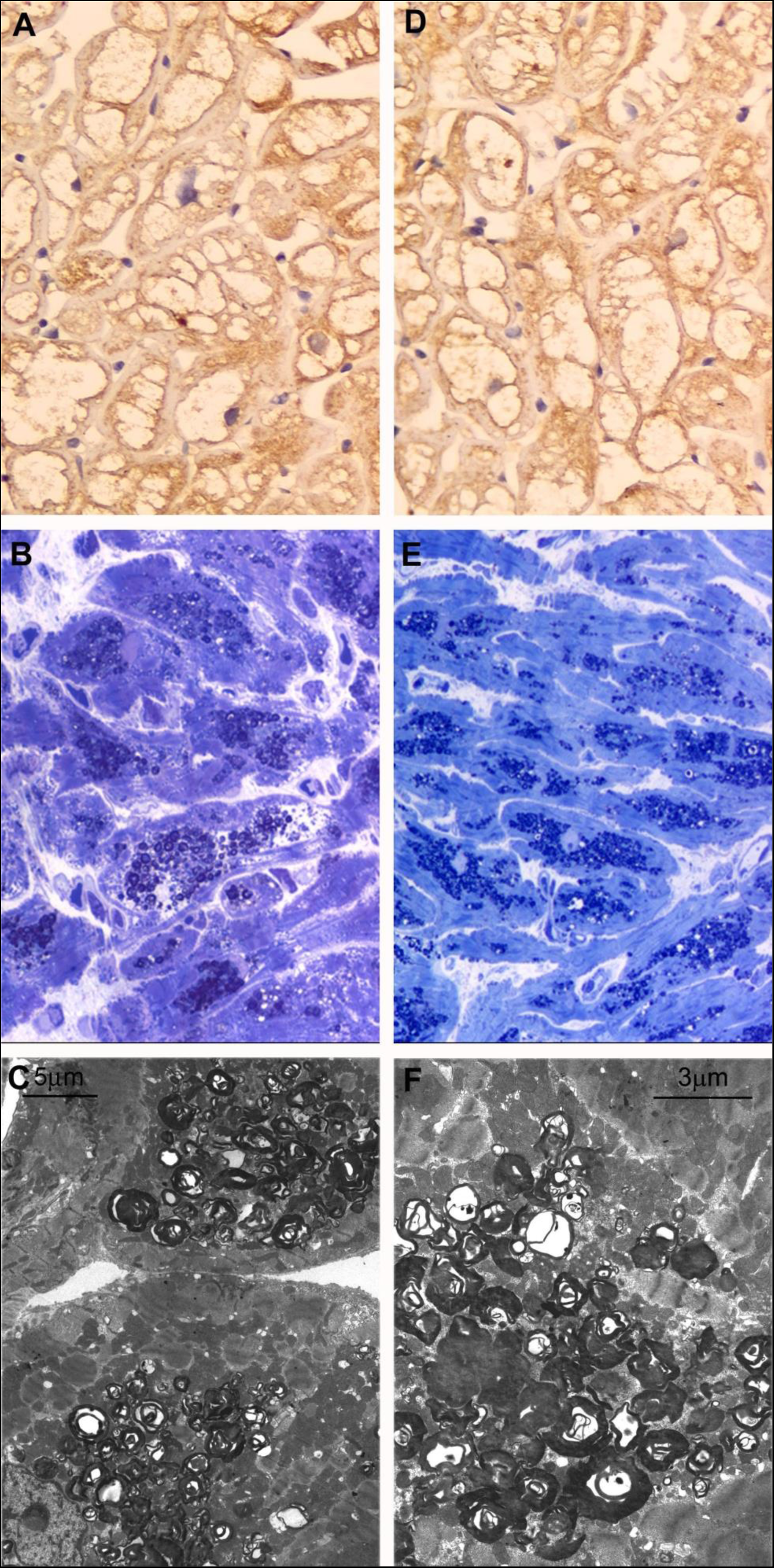
Myocardial immunostaining for alpha-galactosidase A, histologic and ultrastructural features before and 4 hours after enzyme replacement therapy infusion in patients with advanced Fabry Disease Cardiomyopathy. Low cell immunostaining (Panel A, immunoperoxidase for alpha-galactosidase A, 400x) remains unchanged (Panel D), while storage material (Panel B, semithin sections, Azur II staining, 400x; Panel C, transmission electronmicroscopy, uranyl acetate, scale bar=5μm) fails to be removed (Panel E, semithin sections, Azur II staining, 400x; Panel F, transmission electronmicroscopy, uranyl acetate, scale bar=3μm).

**Figure 3.**
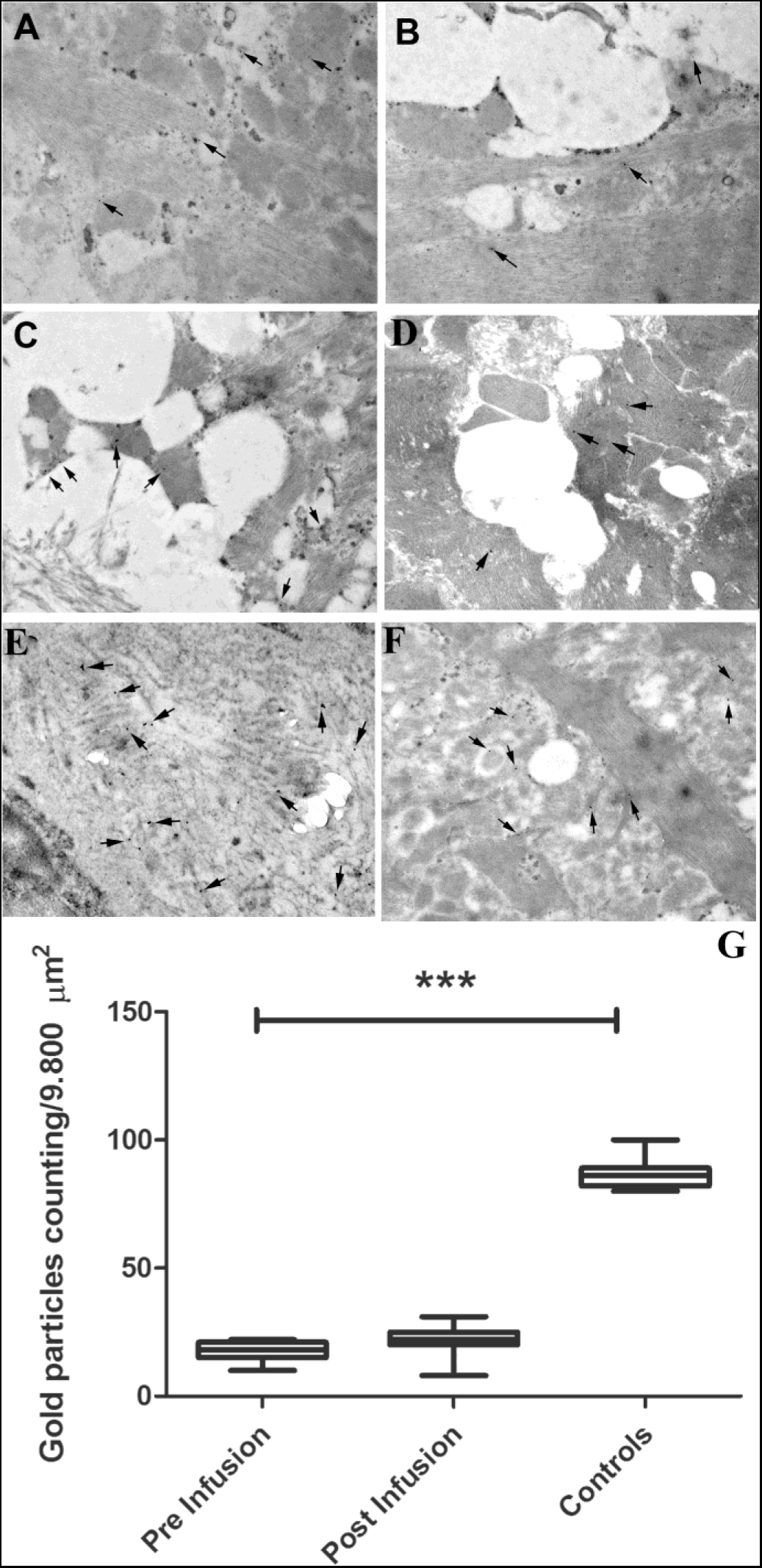
Ultrastructural analysis of cardiac biopsies immunostained with anti-alpha-galactosidase Ab. Arrows indicate the immunogold particles. Panels A-B: Cardiac biopsy from patients in pre-infusion showing only few gold particles (arrows). Panels C-D: Cardiac biopsy from patients in post-infusion .The number of gold particles is mildly (by 1.33 fold) increased (arrows) after 4h ERT infusion. Panels E-F: Cardiac biopsy from healthy donor. Numerous gold particles are visible in the cardiac control tissue (arrows). Original magnifications A-J 20.000 X. Panel G represents in graphic the distribution of gold particles in pre and post agalsidase infusion and the correlation with normal control. p<0.001.

In the 7 patients with histologic evidence of overlapping myocardial inflammation, PCR for the most common cardiotropic was negative.

### Alpha-galactosidase A activity and Immunohistochemistry

Lymphocytic alpha-Gal A activity was reduced in all patients (*Table 1*). Myocardial alpha-galactosidase A activity increased in post infusion samples by about 90% (from 91.6 ± 27.9 to 173.6 ± 47.5 nmol/hr/mg of protein; p<0.001) in the 10 patients with severe FDCM. Immunohistochemistry for the enzyme showed absence of immunostaining in cardiomyocytes of all severe cases (grade 0) in pre-infusion samples (*Table 1 and 2*). In post-infusion samples cardiomyocytes immunostaining mildly improved in some severe cases (*Figure 3, Table 1 and 2*). Immunogold particles increased by 1.33-fold (17.6 ± 3.6 pre infusion vs 21.5 ± 5.9 post) remaining far from normal controls (86.9 ± 6.6).

### Myocardial Mannose-6-phosphate receptors

In patients with severe FDCM the level of M6Pr was significantly lower than controls (4.830 ± 1116,724 vs 14.737± 1145,386; p < 0.001).

## Discussion

The real impact of ERT on FD in general and on FDCM in particular, is still unclear. Results on reduction of myocardial mass are presently controversial and, at least for the advanced form of the disease, rather unsatisfactory ^5–9^.

On the other hand, the poor knowledge on the natural history of the disease in the single patient limits the possibility to evaluate the effect of the treatment even in terms of stabilization of the disease. Finally, uncertainty on ERT efficacy is generated by post-mortem observation of persistent myocardial infiltration after years of ERT administration ^11^ and by experimental studies raising doubts on the ability of the infused enzyme to cross cell membrane ^12^.

In summary there is no definite evidence on the ability of ERT to cross cardiomyocyte membrane, to get activated into the lysosomes, and remove the storage material. On the other hand, mobilization of glycosphingolipids is a key point for reduction of myocardial mass both for lipolysis itself and for inactivation of hypertrophic and proliferative stimuli that appear to be induced by accumulation of GB3 metabolites like lyso-GB3 ^19^.

Our data show that human cardiomyocytes with advanced FDCM are indeed poorly accessible to the usual dosage of ERT. In fact, biopsies after four hours of ERT infusion show low enzyme activity in the myocardium, poor cardiomyocyte immunostaining for alpha-galactosidase A, Immunogold particles increased by only 1.33-fold and no re-absorption of storage material.

It can be argued that time lapse from biopsy studies is too short for these results to be reliable. Indeed, from experimental studies it emerges that the infused enzyme leaves the circulation in one hour and distributes rapidly in peripheral tissues and cell compartment ^20^ reaching its maximal activity in 4 hours ^12^. Then at least the real agalsidase concentration reached in human cardiomyocytes after ERT infusion appear definitely reliable.

Furthermore, it appears of particular interest the analysis of morpho-molecular mechanisms that accompany ERT resistance. Specifically, myocardial fibrosis is increased three folds compared with controls and inflammation overlaps GB3 accumulation in 7 out of 10 patients studied. To this regard, an immune-mediated myocarditis has been recognized in up to 72% of cases in a large histologic study of patients with FDCM as response to highly immunogenic GB3 secretion. ^21^ On the other hand presence of a constitutional secretory pathway for GB3 removal from engulfed cardiomyocytes explain why FDCM patients with absence of alpha-galactosidase A can survive up to 60 years and even longer. Regarding the biological effects of both myocardial fibrosis and inflammation, both these morphologic events cause an expansion of the interstitium and become an obstacle to intracellular ERT internalization ^21^. Furthermore, in the present study, it is reported that the above-mentioned morphologic abnormalities are only an aspect, likely a minor cause, of ERT resistance in FDCM. Indeed, here in are confirmed previous data ^21^ of ERT inefficacy being associated to a severe down-regulation of M6Pr in FDCM (*Figure 4*). In fact, in all patients studied M6Pr appeared reduced by more than 80%. M6Pr are a major molecular complex for uptake and internalization of ERT. Its deficiency may remarkably contribute to a poor ERT impact. Actually, we are unable to explain the origin of this abnormality. In previous studies it was reported a normal expression of mRNA encoding for M6Pr but a 5-fold increase of intracellular ubiquitine suggesting a preserved synthesis but an intracellular enhancement of M6Pr degradation ^22^.

**Figure 4.**
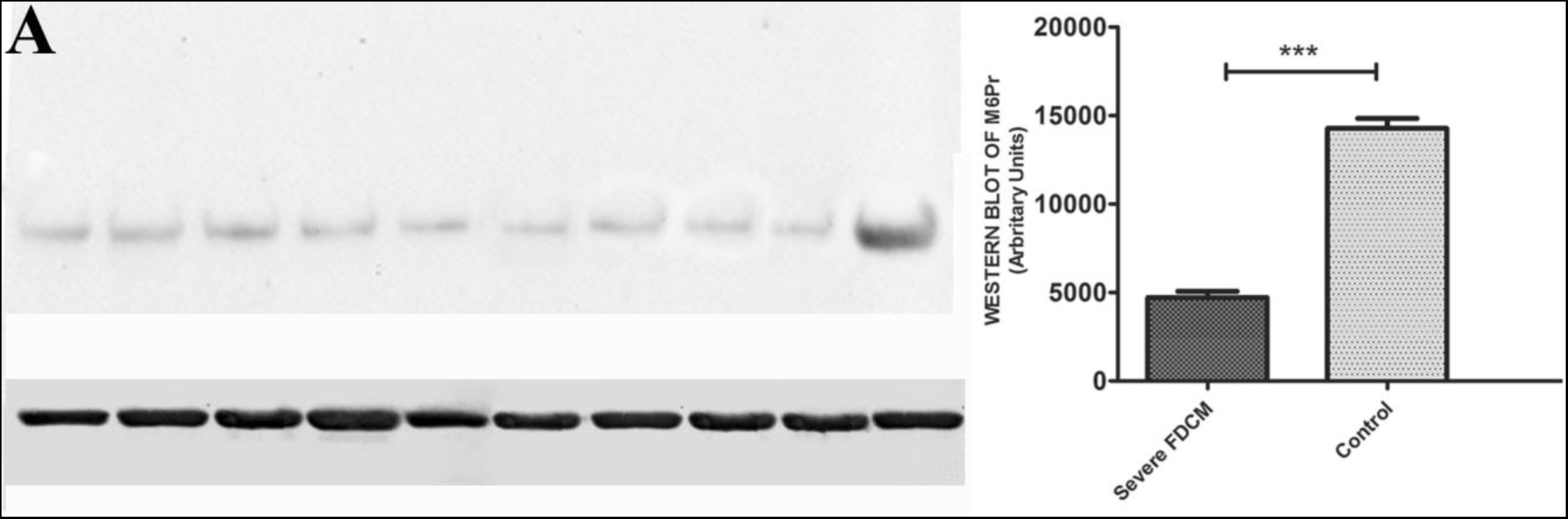
Western blot analysis of all 10 patients of Mannose-6-phosphate Receptor, molecular weight 275 Kd. Panel A Upper: Lane 1 = patient 1; Lane 2 = patient 2; Lane 3 = patient 3; Lane 4 = patient 4; Lane 5 = patient 5; Lane 6 = patient 6; Lane 7 = patient 8; Lane 9 = patient 8. Lane 10 = Control. Alpha sarcomeric actin (43 kDa) was used as a loading control. (Panel A -lower) Panel B: Graphs document western blot of Mannose-6-phosphate Receptor, in all 10 patients with severe FDCM showing 3-fold lower of Mannose-6-phosphate Receptor values in patients versus normal heart (4.830 ± 1116,724 vs 14.737± 1145,386; p < 0.001).

On this base, increase of ERT dosage would be poorly influencing while studies on inhibition of proteasome proteolysis and/or molecular enhancement of M6Pr expression should be taken into consideration. To this regard both Growth hormone and estradiol have been found to increase cellular expression of M6Pr.

## In conclusion

Our study shows a low accessibility to ERT of cardiomyocytes affected by advanced FDCM. It is sustained by myocardial fibrosis, inflammation and severe down-regulation of M6Pr.

## Ethical Committee

The study complies with the Declaration of Helsinki, the locally appointed ethics committee (opinion number 6/2019, La Sapienza University, Rome, Italy) approved the research protocol and informed consent was obtained from all subjects.

## Abbreviations

EMB: Endomyocardial Biopsy
ERT: Enzyme Replacement Therapy
FD: Fabry Disease.
FDCM: Fabry Disease Cardiomyopathy
GB3: Globotriacylceramide.
M6Pr: Mannose-6-phosphate Receptor
TEM: Transmission Electron Microscopy
AB: Antibody

## References.

1. Desnick RJ, Allen KY, Desnick SJ, Raman MK, Bernlohr RW, Krivit W. Fabry’s disease: enzymatic diagnosis of hemizygotes and heterozygotes. Alpha-galactosidase activities in plasma, serum, urine, and leukocytes. J Lab Clin Med. 1973;81:157–171.

2. Nakao S, Takenaka T, Maeda M, Kodama C, Tanaka A, Tahara M, Yoshida A, Kuriyama M, Hayashibe H, Sakuraba H, Tanaka H. An Atypical Variant of Fabry’s Disease in Men with Left Ventricular Hypertrophy. N Engl J Med. 1995;333:288–293.

3. Sachdev B, Takenaka T, Teraguchi H, Tei C, Lee P, McKenna WJ, Elliott PM. Prevalence of Anderson-Fabry Disease in Male Patients With Late Onset Hypertrophic Cardiomyopathy. Circulation. 2002;105:1407–1411.

4. Chimenti C, Frustaci A. Contribution and Risks of Left Ventricular Endomyocardial Biopsy in Patients With Cardiomyopathies: A Retrospective Study Over a 28-Year Period. Circulation. 2013;128:1531–1541.

5. Hughes DA, Elliott PM, Shah J, Zuckerman J, Coghlan G, Brookes J, Mehta AB. Effects of enzyme replacement therapy on the cardiomyopathy of Anderson Fabry disease: a randomised, double-blind, placebo-controlled clinical trial of agalsidase alfa. Heart. 2008;94:153–158.

6. Spinelli L, Pisani A, Sabbatini M, Petretta M, Andreucci M, Procaccini D, Lo Surdo N, Federico S, Cianciaruso B. Enzyme replacement therapy with agalsidase β improves cardiac involvement in Fabry’s disease. Clinical Genetics. 2004;66:158–165.

7. Frustaci A, Chimenti C. Cryptogenic Ventricular Arrhythmias and Sudden Death by Fabry Disease: Prominent Infiltration of Cardiac Conduction Tissue. Circulation [Internet]. 2007 [cited 2023 Dec 16];116. Available from: https://www.ahajournals.org/doi/10.1161/CIRCULATIONAHA.107.723387

8. Koskenvuo JW, Hartiala JJ, Nuutila P, Kalliokoski R, Viikari JS, Engblom E, Penttinen M, Knuuti J, Mononen I, Kantola IM. Twenty-four-month α-galactosidase A replacement therapy in Fabry disease has only minimal effects on symptoms and cardiovascular parameters. J of Inher Metab Disea. 2008;31:432–441.

9. Kovacevic-Preradovic T, Zuber M, Jost CHA, Widmer U, Seifert B, Schulthess G, Fischer A, Jenni R. Anderson-Fabry disease: long-term echocardiographic follow-up under enzyme replacement therapy. European Journal of Echocardiography. 2008;9:729–735.

10. Mehta A, Beck M, Kampmann C, Frustaci A, Germain DP, Pastores GM, Sunder-Plassmann G. Enzyme replacement therapy in Fabry disease: Comparison of agalsidase alfa and agalsidase beta. Molecular Genetics and Metabolism. 2008;95:114–115.

11. Schiffmann R, Rapkiewicz A, Abu-Asab M, Ries M, Askari H, Tsokos M, Quezado M. Pathological findings in a patient with Fabry disease who died after 2.5 years of enzyme replacement. Virchows Arch. 2006;448:337–343.

12. Murray GJ, Anver MR, Kennedy MA, Quirk JM, Schiffmann R. Cellular and tissue distribution of intravenously administered agalsidase alfa. Molecular Genetics and Metabolism. 2007;90:307–312.

13. Keslová-Veselíková J, Hůlková H, Dobrovolný R, Asfaw B, Poupětová H, Berná L, Sikora J, Goláň L, Ledvinová J, Elleder M. Replacement of α-galactosidase A in Fabry disease: effect on fibroblast cultures compared with biopsied tissues of treated patients. Virchows Arch. 2008;452:651–665.

14. Frustaci A, Chimenti C, Ricci R, Natale L, Russo MA, Pieroni M, Eng CM, Desnick RJ. Improvement in Cardiac Function in the Cardiac Variant of Fabry’s Disease with Galactose-Infusion Therapy. N Engl J Med. 2001;345:25–32.

15. Chimenti C, Hamdani N, Boontje NM, DeCobelli F, Esposito A, Bronzwaer JGF, Stienen GJM, Russo MA, Paulus WJ, Frustaci A, Van Der Velden J. Myofilament Degradation and Dysfunction of Human Cardiomyocytes in Fabry Disease. The American Journal of Pathology. 2008;172:1482–1490.

16. Pieroni M, Chimenti C, De Cobelli F, Morgante E, Del Maschio A, Gaudio C, Russo MA, Frustaci A. Fabry’s Disease Cardiomyopathy. Journal of the American College of Cardiology. 2006;47:1663–1671.

17. Aretz HT, Billingham ME, Edwards WD, Factor SM, Fallon JT, Fenoglio JJ, Olsen EG, Schoen FJ. Myocarditis. A histopathologic definition and classification. Am J Cardiovasc Pathol. 1987;1:3–14.

18. Caforio ALP, Pankuweit S, Arbustini E, Basso C, Gimeno-Blanes J, Felix SB, Fu M, Helio T, Heymans S, Jahns R, Klingel K, Linhart A, Maisch B, McKenna W, Mogensen J, Pinto YM, Ristic A, Schultheiss H-P, Seggewiss H, Tavazzi L, Thiene G, Yilmaz A, Charron P, Elliott PM. Current state of knowledge on aetiology, diagnosis, management, and therapy of myocarditis: a position statement of the European Society of Cardiology Working Group on Myocardial and Pericardial Diseases. European Heart Journal. 2013;34:2636–2648.

19. Aerts JM, Groener JE, Kuiper S, Donker-Koopman WE, Strijland A, Ottenhoff R, Van Roomen C, Mirzaian M, Wijburg FA, Linthorst GE, Vedder AC, Rombach SM, Cox-Brinkman J, Somerharju P, Boot RG, Hollak CE, Brady RO, Poorthuis BJ. Elevated globotriaosylsphingosine is a hallmark of Fabry disease. Proc Natl Acad Sci USA. 2008;105:2812–2817.

20. Eng CM, Banikazemi M, Gordon RE, Goldman M, Phelps R, Kim L, Gass A, Winston J, Dikman S, Fallon JT, Brodie S, Stacy CB, Mehta D, Parsons R, Norton K, O’Callaghan M, Desnick RJ. A Phase 1/2 Clinical Trial of Enzyme Replacement in Fabry Disease: Pharmacokinetic, Substrate Clearance, and Safety Studies. The American Journal of Human Genetics. 2001;68:711–722.

21. Frustaci A, Verardo R, Grande C, Galea N, Piselli P, Carbone I, Alfarano M, Russo MA, Chimenti C. Immune-Mediated Myocarditis in Fabry Disease Cardiomyopathy. JAHA. 2018;7:e009052.

22. Frustaci A, Verardo R, Scialla R, Bagnato G, Verardo M, Alfarano M, Russo M. Downregulation of Mannose-6-Phosphate Receptors in Fabry Disease Cardiomyopathy: A Potential Target for Enzyme Therapy Enhancement. JCM. 2022;11:5440.

